# Handling and Packaging of Medical Bags at the Acute Disaster Site Under High Temperature Conditions

**DOI:** 10.1101/2019.12.23.887000

**Authors:** Wataru Ando, Yumika Imamura, Hideyuki Nagashima, Kouji Kondo, Kazunori Nakamura, Katsuya Otori

## Abstract

The handling of medical bags in high temperature disaster sites has not been fully evaluated. In this study, medical bags were assessed in the closed, semi-opened, and opened state, with and without heat insulating material (HIM) and a coolant. The bags were heated for 300 minutes at 35 or 40°C, and the internal temperature was measured. closed and semi-opened bags were able to limit temperature increases in the central part of the bag at both 35 and 40°C to a greater extent than opened bags. When coolant and HIM were used in closed and semi-opened bags, the internal temperature was significantly lower than that seen in the opened state at 40°C (P < 0.01 and P < 0.001, respectively). In conclusion, in high temperature disaster sites, medical bags should be maintained in a semi-opened or closed state using a HIM and coolant.

## Introduction

The medical teams dispatched to the site of a disaster or accident receive specialized training to work in the acute phase where they may be dealing with casualties with multiple injuries. In Japan, the system is based on the disaster medical assistance team (DMAT; [1]. After the large-scale earthquake in 2011 [2], the DMAT was responsible for medical activities during the hyperacute phase, and training of disaster medical teams is ongoing [3]. As these teams will need to administer medicines at the disaster sites, supplies are packaged in medical bags that can be carried to wherever they are needed. The stability of the medicines will differ, and some may be affected by high temperatures or low temperatures, which may reduce their effect [4]. However, drug management at the disaster site has not yet been fully evaluated. The disaster medical team knows empirically that the use of coolants and cold storage boxes is useful for temperature control of pharmaceuticals, but there is no evidence to show any definite measures.

This study aimed to determine the temperature change in medical bags subjected to high temperatures and also to examine the effect of opening the bag and using heat insulating material (HIM) and coolants. The purpose was to propose a method for handling drug bags in high temperature situations.

## Materials and Methods

### Materials

The medical bags used were WJK-1C (Wako Shoji Co., Ltd., Kanagawa, Japan), which were 41 cm long, 49 cm wide, and 23 cm high, with fasteners on three sides. The bag volume was 44 L, and the weight was approximately 2.8 kg. The outer surface was polyvinyl chloride with an inner nylon lining; a 5 mm urethane sheet cushioned the top, sides, and bottom of the bags. Four small nylon bags not containing shock absorbent material were housed inside the bag. Dummy medicines were used in the same quantity, volume, weight, and shape as the 29 standard medicines defined by the DMAT Secretariat of the Ministry of Health, Labor and Welfare of Japan (S1 Table) [5]. The total weight of the packed medical bag was approximately 12 kg, including ampules, soft bags, and packaging. A 10 mm expanded polystyrene foam board (Addtec, Tochigi, Japan) was used as the HIM; 500 g ice packs were used as the coolant (Ice Japan Co., Ltd., Hokkaido, Japan), which were frozen at −20°C before use, and four packs were used per bag. Heating was performed using a convection oven (Espec Corp., Osaka, Japan) as a thermostat box; the temperature was assessed using the MJ-UDL-22 temperature logger (Sato Shoji Inc., Kanagawa, Japan).

### Methods

The bag and dummy drugs were maintained at a constant room temperature of 24.5 ± 1.0°C prior to the study. The temperature logger was placed in the center of the bag, and the dummy drugs and the temperature logger were packed in an identical manner for all experiments. As shown in Fig 1, the bag was set as either 1) completely closed using the fasteners (closed), 2) fasteners open on three sides, with the lid to be partially opened, and the inside of the bag not exposed to the outside (semi-opened), or 3) the inside of the bag fully exposed and the lid released (opened). For each bag status, heat retention was evaluated without HIM and coolant, with HIM only, and with HIM plus coolant.

**Fig 1.**
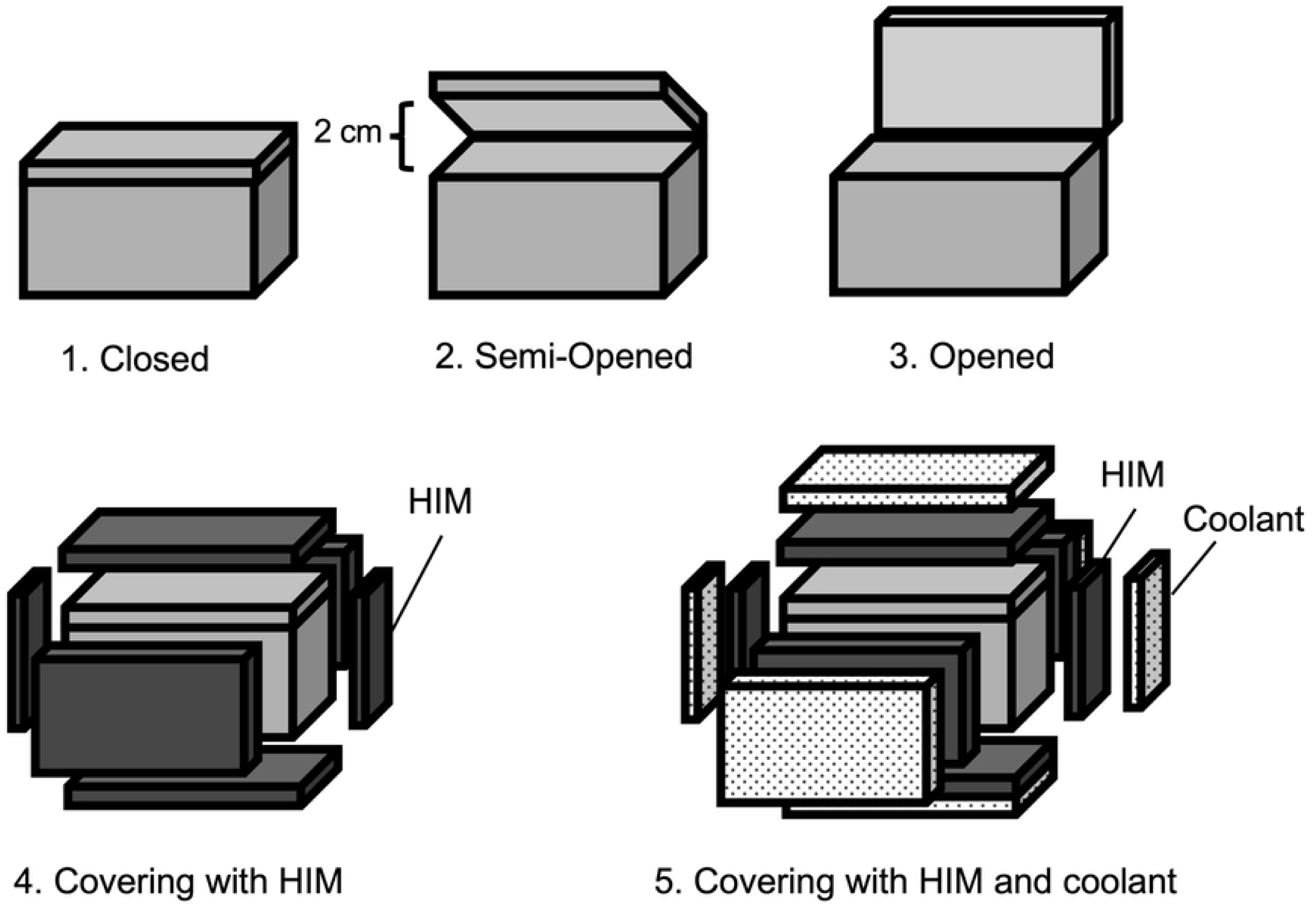
The three different medical bag states used in the experiment.

Initially, the temperature of the thermostat box was set to 35°C or 40°C and stabilized for 1 hour before the experiment. After the bag was placed in the thermostat box, the temperature was measured at the start of the experiment (0) and then at 15, 30, 45, and 60 minutes and every 30 minutes thereafter until 300 minutes. Each experiment was performed in triplicate. The goal was to maintain the temperature in the bag at 15-30°C.

### Statistical analysis

Student’s t test was used to compare the two different temperature groups. For comparison of the three bag status groups, analysis of variance (ANOVA) was used. If the ANOVA was significant, a priori Fisher’s protected least-square difference (LSD) tests were used. The level of statistical significance was p = 0.05.

## Results

### Comparison of the temperature in the medical bags in different states

After 300 minutes at 35 and 40°C, the highest increase in the internal bag temperature was seen in the opened group (Figs 2a and 2b). The temperature did not differ between the closed and semi-opened groups at 35°C and 40°C.

**Figure 2.**
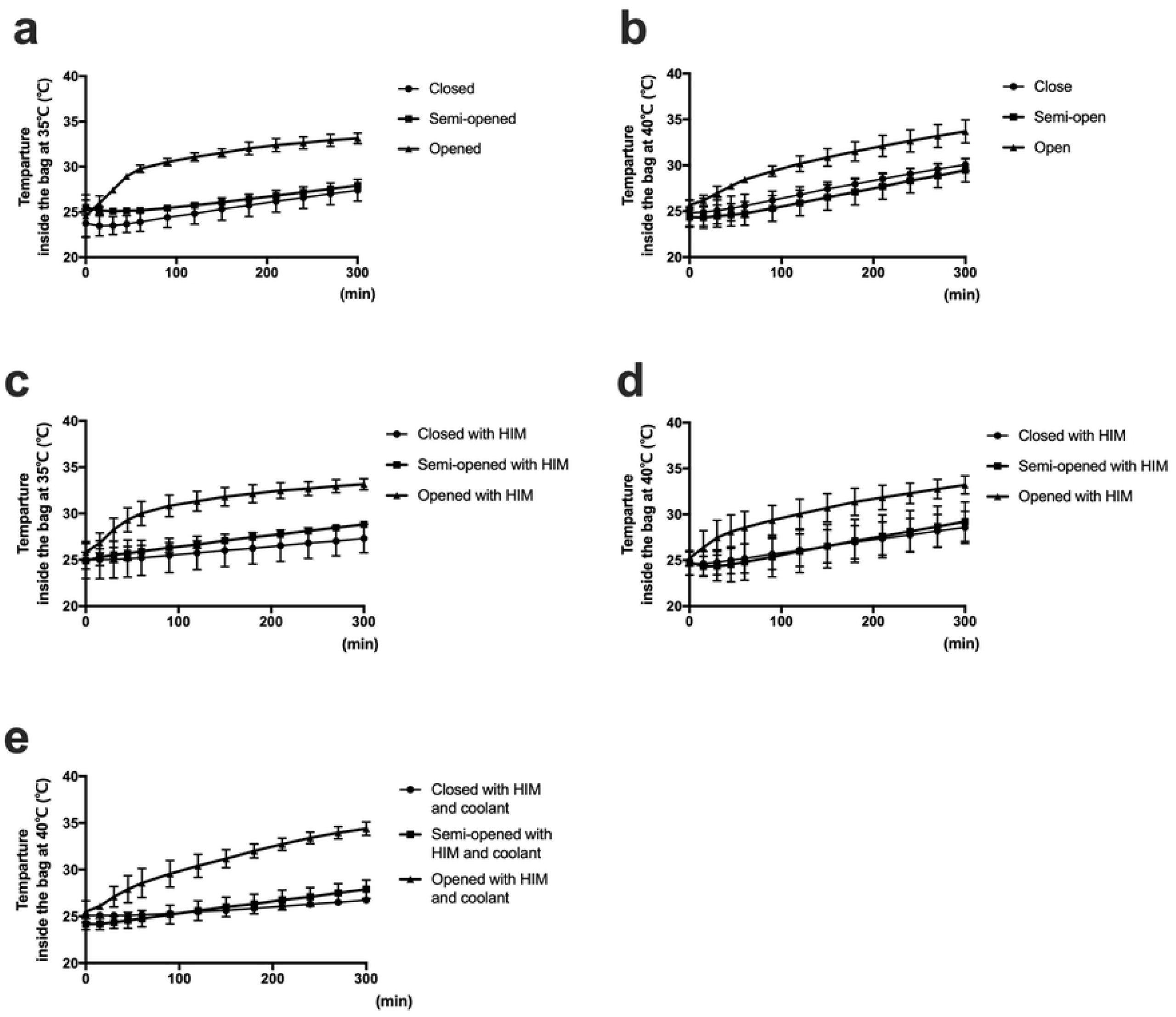
Comparison of temperature changes in medical bags in a closed, semi-opened, and opened state, with or without HIM and coolant. a. 35°C without HIM and coolant, b. 40°C without HIM and coolant, c. 35°C with HIM, d. 40°C with HIM, e. 40°C with HIM and coolant.

### Use of heat insulating materials and coolant

After 300 minutes, the temperature of the bag with HIM was lower than that of the bag without HIM in all bag status groups. At 35°C, the bag with HIM was able to maintain a temperature of < 30°C (Fig 2c). However, at 40°C, the inside temperature exceeded 30°C after 300 minutes (Fig. 2d). At 40°C, the combination of HIM and the coolant maintained the temperature below 30°C in the closed and semi-opened groups, but not in the opened group (Fig 2e).

Fig 3 shows the degree of temperature increase when combining HIM and coolant in each bag state. The temperature increase in the bag was significantly less when using HIM at 35°C and when using both HIM and coolant at 40°C. In particular, in the semi-opened and closed state using HIM at 35°C, the temperature inside the bag was significantly lower than that in the opened state (Fig 3a). The temperature of the semi-opened and closed bags at 40°C was significantly lower than that in the opened state when using HIM and coolant (Fig 3b).

**Figure 3.**
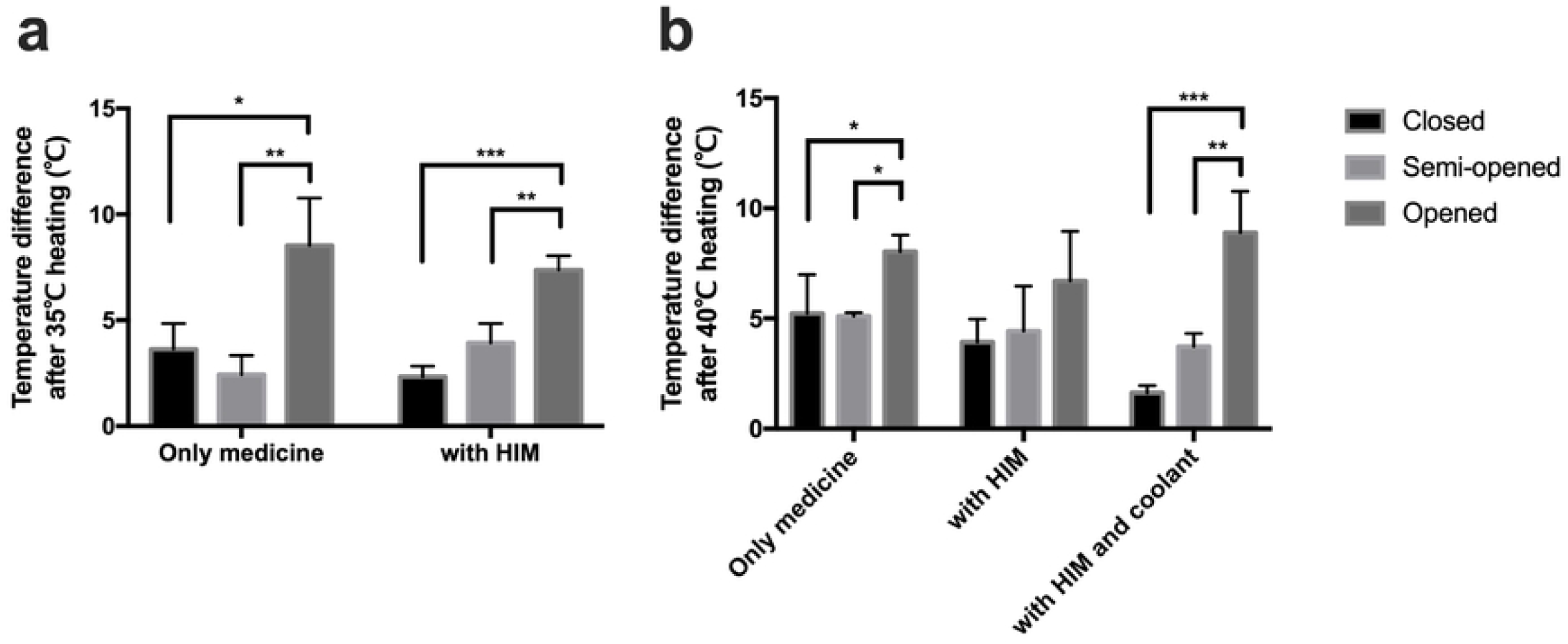
Temperature changes for each bag status after heating. a. heated at 35°C, b. heated at 40°C. ^*^ P < 0.05, ^**^ P < 0.01, ^***^ P < 0.001.

## Discussion

The results of this study show that an opened bag status is more susceptible to the outside temperature than closed and semi-opened bags, and the closed and semi-opened state is more likely to benefit from the use of HIM and coolant.

In this study, a 1 cm-thick expanded polystyrene plate was used as HIM, and it appears that using the thicker boards is more effective. However, because of the reduced internal volume, larger bags may be required, or the number of medications will need to be reduced. While the effectiveness of HIM was not sufficient when the bag was in the opened state, it did provide protection from higher temperatures in the closed or semi-opened state. In addition to polystyrene foam, HIM includes an air cap in which air particles are contained in a resin sheet. The air cap was not assessed in this study because the thickness of this layer changes when stacked. HIM is lightweight and inexpensive and can be readily adapted to the shape of the bag.

While the use of a coolant can limit temperature increases inside the bag, it is important to avoid direct contact between the coolant and the medicine to prevent freezing. In addition, condensation is a consideration as this may encourage the growth of mold. The use of coolant may be insufficient over longer durations.

At the disaster site, the bag would need to be opened when in use, but the current study suggests that a semi-opened state would be preferable to maintain the temperature inside the bag. If the bag is not used for a long time at high temperatures, it should remain closed or semi-opened. As an alternative, drugs that are particularly temperature-sensitive should be placed in the center of the bag. In addition, if a medicine is extremely vulnerable to high temperatures, it may be necessary to manage it separately using a dedicated cooling box and a coolant. There may be differences in stability between original and generic medicines, and drugs included in the DMAT list may differ between teams. Therefore, it will be important to consult a pharmacist about the handling and stability of any medicines used.

Types of heat transfer include direct convection, conduction, and radiation [6]. Heat transfer as conduction and convection affects cold-chain insulated containers with phase change material [7]. In the current study, heat convection is represented by the circulation of heated ambient air. The passage of hot air from outside can be blocked from entering the bag, and convection can be suppressed by packing items evenly in the bag. Direct heat conduction can come from the ground, floor, and walls, as well as the heated environment. In the summer, paved areas of the ground are typically heated by sunlight and the temperature can be higher than that of the environment. Therefore, medical bags should be raised from the ground or flooring. The main source of heat radiation is sunlight, and it is therefore recommended that medical bags be placed in the shade. Particular care will be required for bags stored in stationary cars where the air conditioning will not be operational.

Situations where low temperatures are encountered, i.e., in winter, should also be considered as medicines may freeze if the outside or the ground temperature is below 0°C. There are no reports on how to avoid this situation at disaster sites, but generally a blanket or HIM is used as a countermeasure. An optimal approach to preventing freezing in subzero conditions requires further evaluation.

There are several limitations to the current study. The temperature in a disaster situation is not constant and will fluctuate. Also, the duration of time at 35°C or 40°C may exceed 5 hours, for example, inside a tent. In addition, this study assessed temperature changes on a single day and did not evaluate temperature increases over 2 or more days. Other weather conditions, such as rain, were also not considered. The management and handling of medicines in the DMAT is determined at the discretion of each facility. The list of medicines also varies according to the type and scale of disasters, so the findings of this study may not be applicable to all disaster scenarios. However, quality control of medicines should be conducted as consistently as possible and all staff should be trained in basic medicine management, which should be included more actively in training and operational policies.

## Conclusions

At a high temperature disaster site, the contents of a medical bag can be maintained at a sufficiently low temperature by avoiding the opened state and by keeping the bags semi-opened or closed. The use of HIM and coolant is also advantageous.

## Supporting information

**S1 Table. The based on 29 medicines by the DMAT Secretariat of the Ministry of Health, Labor and Welfare of Japan**.

